# Anti-SARS-CoV-2 hyperimmune immunoglobulin provides potent and robust neutralization capacity and antibody-dependent cellular cytotoxicity and phagocytosis induction through N and S proteins

**DOI:** 10.1101/2021.06.11.447942

**Authors:** José María Díez, Carolina Romero, María Cruz, Peter Vandeberg, W. Keither Merritt, Edwards Pradenas, Benjamin Trinité, Julià Blanco, Bonaventura Clotet, Todd Willis, Rodrigo Gajardo

## Abstract

**Background:** Although progressive COVID-19 vaccinations provide a significant reduction of infection rate in the short-to mid-term, effective COVID-19 treatments will continue to be an urgent need.

**Methods:** We have functionally characterized the anti-SARS-CoV-2 hyperimmune immunoglobulin (hIG) prepared from human COVID-19 convalescent plasma. SARS-CoV-2 virus neutralization was evaluated by four different methodologies (plaque reduction, virus induced cytotoxicity, TCID50 reduction and immunofluorimetry-based methodology) performed at four different laboratories and using four geographically different SARS-CoV-2 isolates (one each from USA and Italy; two from Spain). Two of the isolates contained the D614G mutation. Neutralization capacity against the original Wuhan SARS-CoV-2 straom and variants (D614G mutant, B.1.1.7, P.1 and B.1.351 variants) was evaluated using a pseudovirus platform expressing the corresponding spike (S) protein. The capacity to induce antibody-dependent cellular cytotoxicity (ADCC) and antibody-dependent cellular phagocytosis (ADCP) was also evaluated.

**Results:** All the SARS-CoV-2 isolates tested were effectively neutralized by hIG solutions. This was confirmed by all four methodologies showing potent neutralization capacity. Wild-type SARS-CoV-2 and variants were effectively neutralized as demonstrated using the pseudovirus platform. The hIG solutions had the capacity to induce ADCC and ADCP against SARS-CoV-2 N and S proteins but not the E protein. Under our experimental conditions, very low concentrations (25-100 µg IgG/mL) were required to induce both effects. Besides the S protein, we observed a clear and potent effect triggered by antibodies in the hIG solutions against the SARS-CoV-2 N protein.

**Conclusions:** These results show that, beyond neutralization, other IgG Fc-dependent pathways may play a role in the protection from and/or resolution of SARS-CoV-2 infection when using hIG COVID-19 products. This could be especially relevant for the treatment of more neutralization resistant SARS-CoV-2 variants of concern.

## 1. Introduction

One year after the WHO declared the respiratory disease, COVID-19, a pandemic (March 2020), the global situation is still of great concern as new cases continue to rise [1]. Although several vaccines have been marketed and others are in different stages of development [2], limited supplies makes their immediate and wide availability unlikely. Therefore, the vaccination of the worldwide population progresses slowly [3]. Even with an expectation of a reduction of the infection rate in the short-to mid-term, infected patients remain in need of effective treatments.

Currently, there is no effective standardized treatment for COVID-19, although multiple therapeutic options are available to tackle the different symptoms of the disease or to directly target SARS-CoV-2, its genome and cell cycle [4]. Among the available therapeutic strategies, passive immunization using COVID-19 convalescent plasma (CCP) or hyperimmune globulin (hIG: IgG enriched with anti-SARS-CoV-2 antibodies) is of particular relevance [5]. As the SARS-CoV-2 pandemic spreads and vaccination of the general population progresses, commercial IgG medicinal products derived from healthy plasma donors become gradually enriched in anti-SARS-CoV-2 antibodies [6]. To date, the antibody levels in the general population are still low [6], therefore the plasma collected, and the final product produced cannot yet be considered a hyperimmune.

hIG is typically prepared from pools of 100–1000 liters from CCP donors. hIG products have a high titer of neutralizing antibodies against SARS-CoV-2 in a standardized and concentrated product [7]. This represents a relevant advantage over treatment with CCP. Key targets of anti-SARS-CoV-2 antibodies include: the S protein [8], which is responsible for viral entry through recognition of the primary host cellular receptor angiotensin-converting enzyme-2 (ACE2); and the N protein [9], which makes up the helical nucleocapsid. The E protein, which is a small polypeptide, and the M protein, which is embedded in the envelope [10], have been less studied as potential immune targets.

Importantly, IgG possess other antiviral properties, beyond neutralization, which have been described for CCP, including antigen-dependent Fc functions, e.g. antibody dependent cellular phagocytosis (ADCP) [11], antibody dependent cellular cytotoxicity (ADCC) [12], complement-mediated cytotoxicity [13], etc. These well-described effector functions of antibodies (mediated by the interaction of immunoglobulin Fc with cellular Fc receptors) may add to neutralizing activity and may enable non-neutralizing antibodies or antibodies with poor-neutralizing capacity to block or clear infection. hIG efficacy is being tested in ongoing randomized clinical trials in inpatients (IV administration) [14], and outpatients (SC administration) [15].

Apart from their SARS-CoV-2 neutralizing capacity, no further antiviral capacity of hIG has been experimentally demonstrated. In this study, we report an extensive functional characterization of an hIG product [7]. We performed neutralization assays on several virus isolates and on a pseudovirus expressing the most relevant S variants to date, and, for the first time, we evaluated the capacity of hIG to trigger antigen-dependent IgG Fc functions.

## 2. Materials and Methods

### 2.1 Study Design

The anti-SARS-CoV-2 hyperimmune globulin 10% (hIG) (Grifols, Barcelona, Spain) prepared from CCP was functionally characterized in vitro. In the first series of studies, neutralization of four geographically diverse isolates of SARS-CoV-2 were assessed by four different methodologies (plaque reduction, protection from virus-induced cytotoxicity, TCID50 reduction and immunofluorimetry-based methodology) at four different laboratories. Secondly, the capacity of the hIG to neutralize SARS-CoV-2 variants was evaluated using a pseudovirus test platform expressing the S proteins of the relevant variants. Finally, the capacity of hIG to induce antibody-dependent cellular cytotoxicity (ADCC) and antibody-dependent cellular phagocytosis (ADCP) on the same samples and the viral protein responsible of eliciting these responses were evaluated. Positive (CCP) and negative single-donation plasmas were used for comparison.

### 2.2 Cell Lines and Culture

Each of the four testing laboratories used their own cell lines, as follows:

At NIH, Vero cells were acquired from the American Type Culture Collection (ATCC #CCL-81; Manassas, VA, USA) and cultured in DMEM (Lonza, Walkersville, MD, USA) containing 2% FBS (SAFC Biosciences, Lenexa, KS, USA).

At CReSA-IRTA, Vero cells were obtained from the ATCC (ATCC CRL-1586) and cultured in DMEM (Lonza, Basel, Switzerland) supplemented with 5% FBS (EuroClone, Pero, Italy), 100 U/ml penicillin, 100 μg/mL streptomycin and 2 mM glutamine (all from ThermoFisher Scientific, MA, USA). In the post infection medium, the percentage of FBS was 2%.

At CNB-CSIC, Vero cell lines were kindly provided by Dr E Snjider (University of Leiden Medical Center, The Netherlands). Vero cell lines were cultured in Dulbecco’s modified Eagle medium (DMEM) supplemented with 25 mM HEPES buffer, 2 mM glutamine (Sigma-Aldrich, MI, USA), 1% nonessential amino acids (Sigma-Aldrich), 10% fetal bovine serum (FBS; BioWhittaker, Inc., MD, USA). In the post infection semisolid medium, the percentage of FBS was reduced to 2%, and diethylaminoethyl (DEAE)-dextran (Sigma-Aldrich) was added to a final concentration of 0.08 mg/mL.

At Texcell, Vero cells (provided by Pasteur Institut) were cultured in DMEM (Sigma ref: D6546) 4% containing FBS (VWR Ref: 97068-085) 2% Glutamine (Lonza Ref: BE17-605E). Inoculation medium contained DMEM 2% FBS 1% Glutamine.

At IrsiCaixa, HEK 293T cells overexpressing WT human ACE-2 (Integral Molecular, USA) were used for pseudovirus neutralization assays.

### 2.3 SARS-CoV-2 Strains

Stock viruses were prepared by collecting the supernatant from Vero cells, as previously described [16]. Each neutralization methodology and four laboratories used their own SARS-CoV-2 strain as follows:

For CBIFA at NIH, SARS-CoV-2 (GenBank #MT020880) was provided by the U.S. Centers for Disease Control and Prevention (Washington isolate, CDC; Atlanta, GA, USA). It was isolated from the first US COVID-19 patient [17].

For CCLA at IRTA-CReSA, SARS-CoV-2 isolated from nasopharyngeal swab from an 89-year-old male patient from Badalona (Spain) in March 2020 (accession ID EPI ISL 418268 at GISAID repository: http://gisaid.org) with the following mutations Spike D614G, NSP12 P323L.

For PFU-based neutralization at CNB-CSIC, SARS-CoV-2MAD6 isolated from nasopharyngeal swab from a 69-year-old male patient from Hospital “12 de Octubre” in Madrid. The virus was isolated, plaque cloned three times and amplified in Coronavirus laboratory at CNB-CSIC. Full-length virus genome was sequenced and it was found to be identical to SARS-CoV-2 reference sequence (Wuhan-Hu-1 isolate, GenBank MN908947), except for the presence of a silent mutation C3037>T, and two mutations leading to aa changes: C14408>T (in nsp12) and A23403>G (D614G in S protein).

For TCID50 based at Texcell, 2019-nCoV strain 2019-nCoV/Italy-INMI1 (https://www.ncbi.nlm.nih.gov/nuccore/MT066156) was used. It was isolated from the first case of COVID-19 in Italy [18].

### 2.4 Convalescent Plasmas

SARS-CoV-2 antibody-positive plasmas were collected by plasmapheresis from CCP donors (single donation) at US plasma collection centers within the Grifols network (Biomat USA Inc., Interstate Blood Bank Inc., Talecris Plasma Resources, Inc.). CCP was collected during the first half 2020. The CCP donors had COVID-19 with different degrees of severity from mild to hospitalized.

COVID-19 specific antibody levels of the plasma units were classified as high (positive at least at 1/10000 dilution), medium (positive at 1/1000), and low (positive at 1/100) as determined by anti-SARS-CoV-2 S ELISA methods: human anti-SARS-CoV-2 virus spike 1 [S1] IgG ELISA Kit (Alpha Diagnostic Intl., Inc.), against S1 subunit spike protein; EI-2606-9601-G, Anti-SARS-CoV-2 IgG ELISA Kit (Euroimmun AG, Luebeck, Germany), against structural protein (S1 domain); DEIASL019, SARS-CoV-2 IgG ELISA Kit (Creative Diagnostics), against virus lysate.

SARS-CoV-2 antibody-negative plasma (pre-pandemic collection during 2019) was used as a negative control.

### 2.5 Experimental Product

The hIG used in this study is a highly purified IgG product manufactured through caprylate/chromatography purification process using CCP. Manufacturing CCP pools were constituted of around 100 different plasma donations. In the hIG manufacturing process, IgG concentration and the specific antibody to SARS-CoV-2 is increased over 10-fold from CCP to the final product. Protein content is 100% IgG, consisting of 98% monomer and dimer forms. Potentially accompanying impurity proteins (IgM, IgA, and anti-A, anti-B and anti-D antibodies) are reduced to minimal levels. Further details of hIG characteristics are available elsewhere [7].

### 2.6 SARS-CoV-2 Neutralization Experiments

The following infectivity neutralization methodology were used:

A cell-based immunofluorescence assay (CBIFA) was used at NIH. This assay quantifies the anti-SARS-CoV-2 neutralization titer using a dilution series of test material tested for inhibition of infection of cultured Vero E6 cells by SARS-CoV-2 at a multiplicity of infection of 0.5. Potency was assessed using a cell-based immunosorbent assay to quantify infection by detecting the SARS-CoV-2 nucleoprotein using a specific antibody raised against this protein (Sino Biological, Wayne, PA). The secondary antibody (Life Technologies, San Diego, CA) was conjugated to Alexa Fluor 594 and cells were quantified with an Operetta high content imaging system (Perkin Elmer, Waltham, MA). A minimum of 16,000 cells are counted per sample dilution across four wells - two each in duplicate plates. Data are reported based on a 4-parameter regression curve (using a constrained fit) as a 50% neutralization titer (half maximal inhibitory dilution : ID50) [7, 19].

A cytopathic-cytotoxicity luminometry assay (CCLA) was used at IRTA-CReSA. This neutralization assay measured the cytopathic/cytotoxic virus-induced effect by detecting cellular enzymatic activity after incubation with a given amount of the relevant virus and comparing this with the relevant untreated control. For this assay, a fixed concentration of the SARS-CoV-2 stock (101.8 TCID50/mL, a concentration that achieves 50% cytopathic effect) was mixed with decreasing concentrations of the hIG samples and added to Vero E6 cells. To assess potential plasma-induced cytotoxicity, Vero E6 cells were also cultured with the same decreasing concentrations of plasma in the absence of SARS-CoV-2. Cytopathic or cytotoxic effects of the virus or plasma samples were measured at 3 days post-infection, using the Cell Titer-Glo luminescent cell viability assay (Promega, WI, USA). Luminescence was measured in a Fluoroskan Ascent FL luminometer (Thermo Fisher Scientific). ID50 values were determined from the fitted neutralization curves as the plasma dilutions that produced 50% neutralization. Details of the technique are available elsewhere [20, 21].

Plaque forming units (PFU)-based neutralization assay was used at CNB-CSIC. This neutralization assay was based on the reduction in PFU after exposing a given amount of virus to the product to be characterized and comparing with the untreated control. hIG samples were serially diluted in Dulbecco’s phosphate-buffered saline (Gibco, ThermoFisher Scientific, MA, USA) and incubated with 300 PFUs of SARS-CoV-2. Aliquots of each hIG dilution-virus complex were added in duplicate to confluent monolayers of Vero E6 cells. The neutralization potency of the hIG product (ID50 value) was expressed as plaque reduction neutralization test (PRNT50) value, calculated as the -log10 of the reciprocal of the highest hIG dilution to reduce the number of plaques by 50% compared with the number of plaques without IVIG [21].

Median Tissue Culture Infectious Dose (TCID50)-based microneutralization assay was used at Texcell. Briefly, immunoglobulins dilutions are incubated with 400 TCID50 (equivalent to 277 viral infectious units) for 1h at 37° C and 5% CO2. After that a microplate seeded before with Vero cells is inoculated adding 35 µL/well inmmunoglobulin/virus mixture, this mixture is absorbed for 1 h at 37° C in 5% CO2 atmosphere and after that 65 µL cell culture medium overlay is added. The plate is incubated for 6 days at 37° C in 5% CO2, then the plate is read for the presence or absence of cythopathic effect (CPE). The viral titer is expressed in “dose infecting 50% of tissue cultures per mL” with a confidence interval of 95%. For neutralization plate, the ID50 value was expressed as the Neutralization Titer 50 (NT50) value, calculated as the antibody titer neutralizing sample according to the Spearman-Kärber formula. The NT50 corresponds to the dilution of sample which prevent the cells from CPE (no lysis) in 50% of the replicates. The criteria for the validation of the run were: back titration of the virus in the TCID50 criteria; integrity of the uninfected cells (medium control only); and absence of cell layer or presence of CPE in infected wells (virus control only).

### 2.7 Generation of Spike Expression Plasmids

SARS-CoV-2.SctΔ19 Wuhan, B.1.1.7, P.1 and B.1.351 variants were generated (GeneArt) from the full protein sequence of the respective spike sequences, with a deletion of the last 19 amino acids in C-terminal [22]. Sequences were human-codon optimized and inserted into pcDNA3.1(+). The G614 spike mutant was generated by site-directed mutagenesis as previously described [23]. In brief, SARS-COV-2.SctΔ19 Wuhan plasmid was amplified by PCR with Phusion DNA polymerase (Thermo Scientific, ref F-549S) and the following primers: 5’-TACCAGGgCGTGAACTGTACCGAAGTGCC-3’ and 5’-GTTCACGcCCTGGTACAGCACTGCCAC-3’. PCR was 20 cycles with an annealing temperature of 60° C and an elongation temperature of 72° C. PCR product was then treated for 3 hours with the DpnI restriction enzyme (Thermo Scientific, ref 00909083), to eliminate template DNA, and transformed into supercompetent E. coli. Final mutated DNA was then fully sequenced for validation.

### 2.8 Pseudovirus Generation and Neutralization Assay

In these experiments performed at IrsiCaixa AIDS Research Institute, HIV reporter pseudovirus expressing SARS-CoV-2 S protein and Luciferase were generated using a plasmid coding for a non-replicative HIV reporter pNL4-3.Luc.R-.E-obtained from the NIH AIDS Reagent Program [24] and the spike expression plasmids (described above). Expi293F cells were transfected using ExpiFectamine293 Reagent (Thermo Fisher Scientific) with pNL4-3.Luc.R-.E- and SARS-CoV-2.SctΔ19 (Wuhan, G614, B.1.1.7, P1 and B.1.351), at an 8:1 ratio, respectively. Control pseudovirus was obtained by replacing the S protein expression plasmid with a VSV-G protein expression plasmid as reported [25]. Supernatants were harvested 48 hours after transfection, filtered at 0.45 µm, frozen, and titrated on HEK293T cells overexpressing WT human ACE-2 (Integral Molecular, USA). This neutralization assay has been previously validated in a large subset of samples [26].

Neutralization assays were performed in duplicate as previously described [26]. Briefly, in Nunc 96-well cell culture plates (Thermo Fisher Scientific), 200 TCID50 of pseudovirus was preincubated with three-fold serial dilutions of samples for 1 hour at 37° C. Then, 1×104 HEK293T/hACE2 cells treated with DEAE-Dextran (Sigma-Aldrich) were added. Results were read after 48 hours using the EnSight Multimode Plate Reader and BriteLite Plus Luciferase reagent (PerkinElmer, USA). The values were normalized, and the ID50 (the reciprocal dilution inhibiting 50% of the infection) was calculated by plotting and fitting the log of plasma dilution vs. response to a 4-parameters equation in Prism 8.4.3 (GraphPad Software, USA).

### 2.9 ADCC-ADCP Induction Experiments

p Antibody-dependent cell-mediated cytotoxicity (ADCC) and antibody-dependent cellular phagocytosis (ADCP) are mechanisms through which virus-infected or other diseased cells are targeted for destruction or elimination by multiple components of the cell-mediated immune system. These specific mechanisms were assayed using bioluminescent reporter assays for quantifying ADCC/ADCP pathway activation by several therapeutic antibody drugs.

ADCC and ADCP techniques were evaluated using the ADCC Reporter Bioassay, Core Kit, Promega (ADCC Reporter Bioassays, FcγRIIIa V158 variant (high affinity), Catalog number G7010, G7018, Promega Corporation) and FcγRIIa-H (high affinity) ADCP Bioassay, Core kit, Promega (ADCP FcγRIIa-H Reporter Bioassay, Core Kit, Promega Corporation Catalog number: G9995) respectively, following the manufacturer’s guidelines.

Briefly, in a first set of experiments, SARS-CoV-2 antigens (N, S and E proteins) were used for coating 96-well plates, and several sample types were evaluated for ADCC and ADCP functionalities using the corresponding Reporter Bioassay kit (Promega Corporation). ADCC and ADCP induction was expressed as induction ratio (IR), which corresponds to the detected signal versus the 1U/mL kit calibrator.

Additionally, the antigen coating capacity was evaluated and confirmed by SARS-CoV-2 ELISAs (Alpha Diagnostic Human anti-SARS-CoV-2 S1 IgG ELISA, RV-405200; and Alpha Diagnostic Human anti-SARS-CoV-2 Nucleoprotein IgG ELISA, RV-405100) using Corning® 96-well Flat Clear Bottom White Polystyrene TC-treated Microplates (Corning Ref. 3903), the three SARS-CoV-2 antigens (N, S and E), and three different sample types (prepandemic IgG within Gamunex C, and IgG ELISA SARS-CoV-2 high positive and negative single-donation plasmas).

SARS-CoV-2 spike glycoprotein expressing HEK293T cells (Innoprot, Reference: P30908) were then used to verify these functionalities (ADCC/ADCP) for SARS-CoV-2 S antigen. A SARS-CoV-2 spike glycoprotein cell line was stably developed transfecting the HEK293T cell line with a SARS-CoV-2 spike glycoprotein expression plasmid (Innoprot, Derio, Basque Country, Spain). This cell line provides consistent levels of expression of human SARS-CoV-2 spike glycoprotein on the cell surface.

In these experiments, samples were assayed at increasing concentrations in order to perform a kinetic curve (concentration/response curve) for the dynamic evaluation of ADCC and ADCP functionalities in all sample types.

Plasma samples with high, medium, low, and null positivity for COVID-19 infection (determined by anti-SARS-CoV-2 S ELISA methods) were used as comparators.

### 2.10 Statistics

Neutralization titers were calculated using GraphPad Prism 8 version 8.4.3. nonlinear regression curve fit as half-maximal inhibitory dilution (ID50). The titers obtained for different batches are expressed as the mean value ± standard deviation (SD).

## 3. Results

### 3.1 SARS-CoV-2 Neutralization

e All the methods tested demonstrated neutralization of infectivity by hIG in the four SARS-CoV-2 isolates (one from USA, one from Italy, and two from Spain). ID50 results are shown in Table 1. Differences in ID50 are ascribed to differences in the methodologies employed reflecting their differential sensitivities. The D614G mutation was present in the isolates from Spain.

**Table 1.**
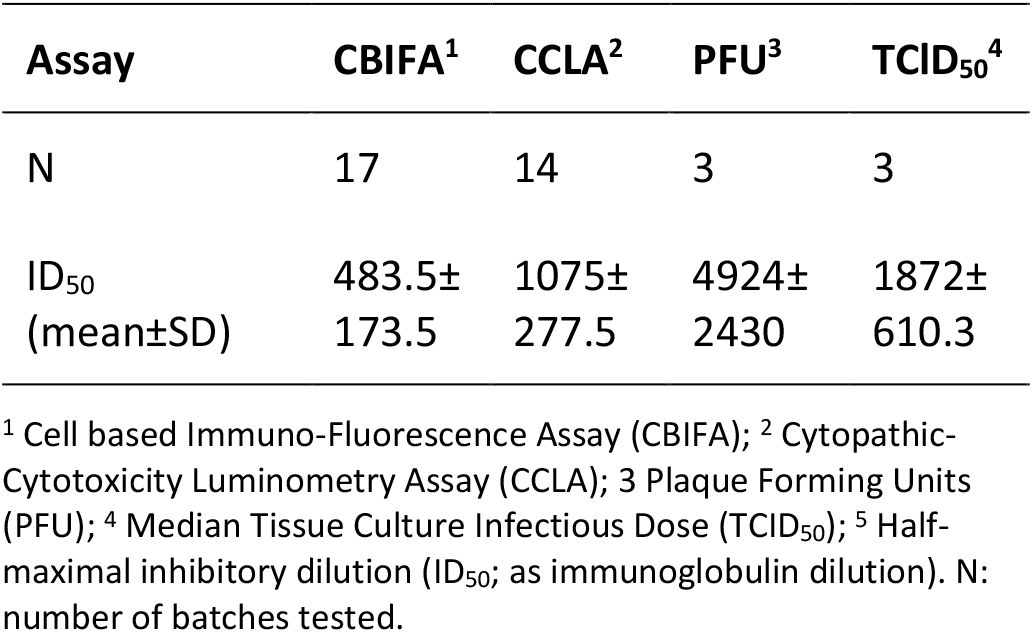
SARS-CoV-2 infectivity neutralization by hyperimmune IG (half-maximal inhibitory dilution, ID_50_)

### 3.2 SARS-CoV-2 Variants Pseudovirus Neutralization

The neutralization assays with pseudovirus demonstrated the neutralization capability against wild-type (original Wuhan virus) and all the variants: D614G, B.1.1.7 UK, P.1 Brazilian and B.1.351 South African (Table 2 and Figure 1). The levels of neutralization were very similar for Wuhan virus and the D614G and B.1.1.7 spikes and lower for the P1 and B.1.351 spikes, but still showed consistent neutralization capacity. As a negative control a normal IgG IVIG (pre-pandemic) was used showing no detectable neutralization with this pseudovirus platform.

**Table 2.**
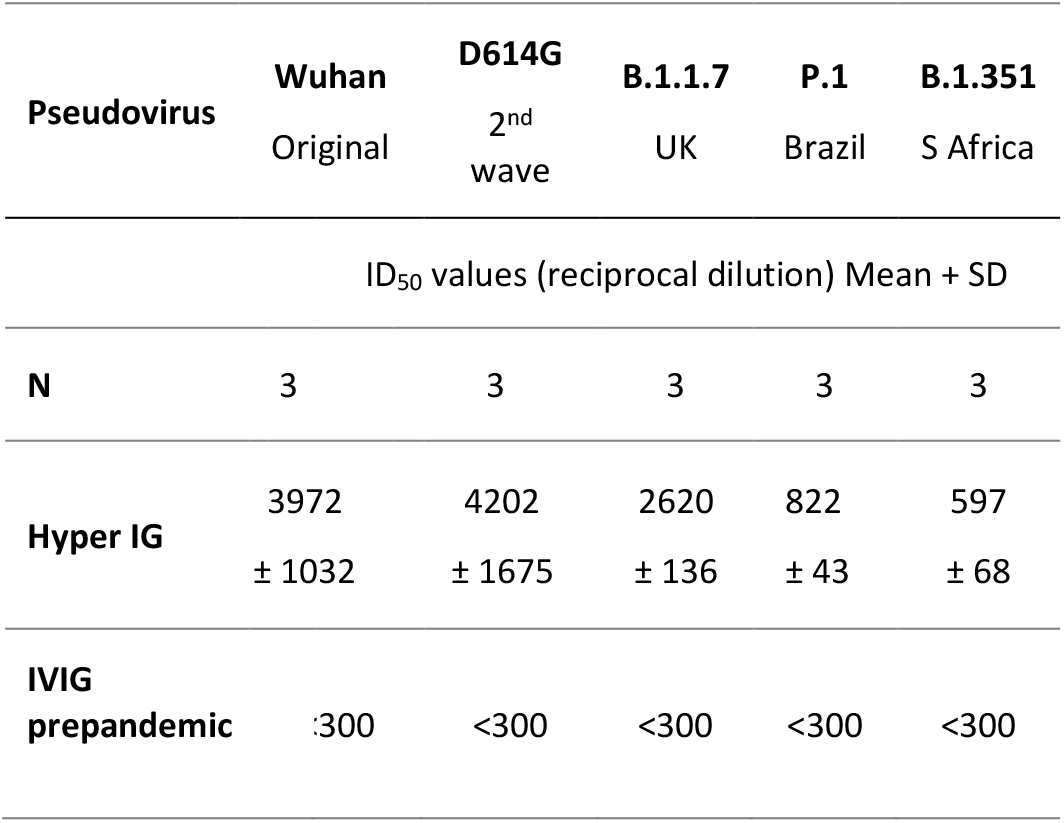
SARS-CoV-2 variants infectivity neutralization by hyperimmune IG (half-maximal inhibitory dilution, ID_50_)

**Figure 1.**
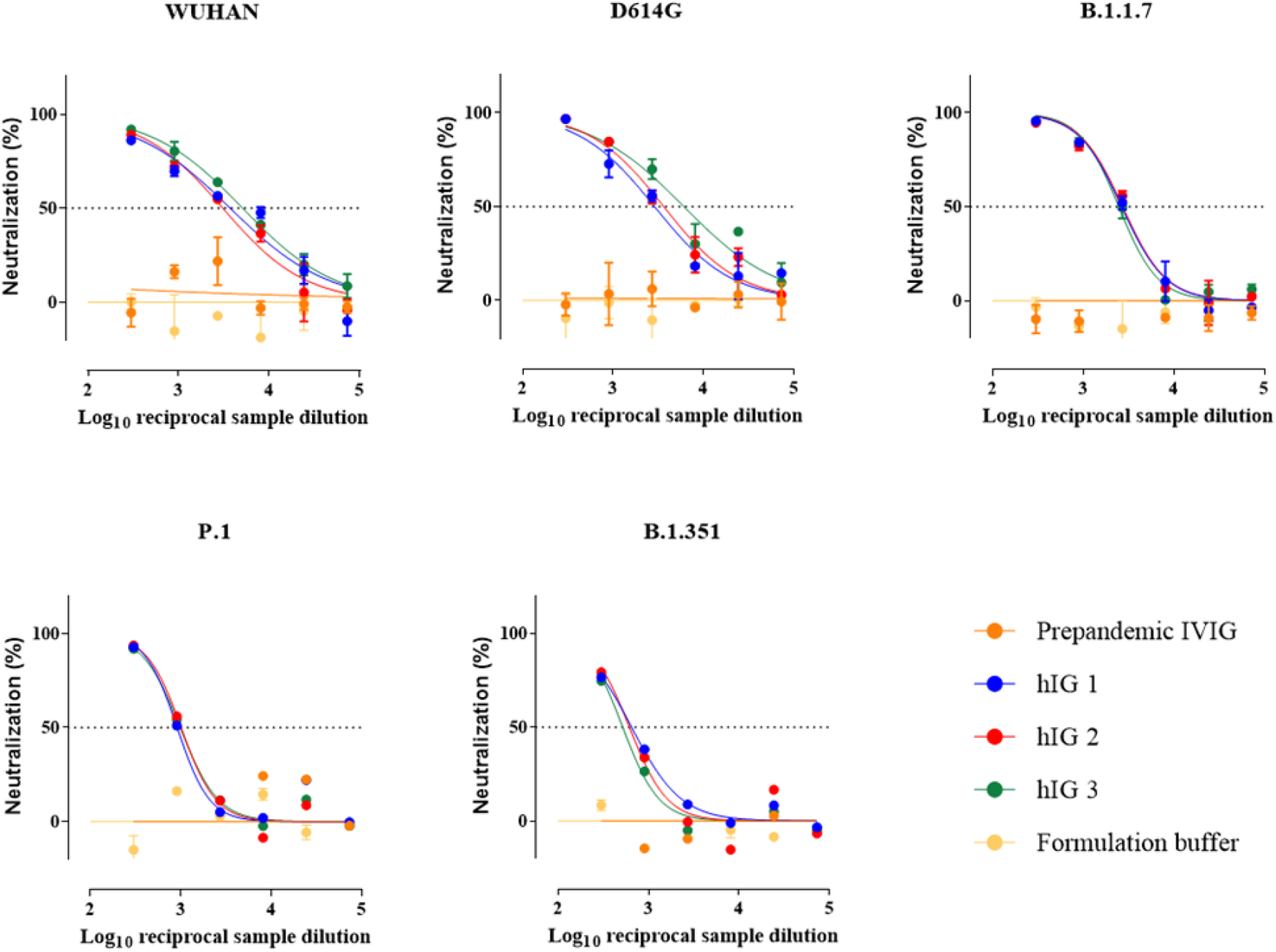
SARS-CoV-2 variants infectivity neutralization by hyperimmune IG. Neutralization curves for the indicated SARS-CoV-2 variants (pseudovirus expressing S protein). A prepandemic IGIV and the formulation buffer were tested in parallel as negative controls.

### 3.3 ADCC-ADCP Induction

A strong ADCC and ADCP induction ratio (IR) by hIG was observed on plates coated with SARS-CoV-2 N antigen (IR of 6 or higher for ADCC and a IR of 10 or higher for ADCP), at low hIG concentrations (µg IgG/mL) but not with the E and S antigens. Some ADCP induction by the S antigen was observed at higher concentrations of hIG (IR around 2 at 5 mg IgG/mL) (Supplementary information Figures S1 and S2). Similarly, ADCC and ADCP induction response for pre-pandemic plasma and pre-pandemic IVIG samples (negative controls) were very weak as expected (IR around 1) at any concentration. These results are summarized in Figures 2a and 2b.

**Figure 2A.**
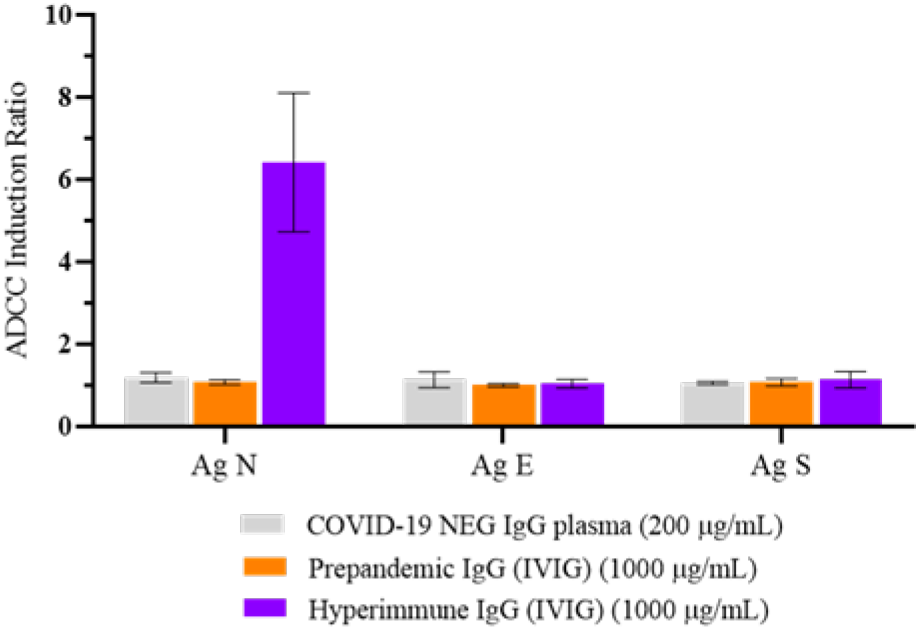
ADCC induction ratio with coated SARS-CoV-2 antigens. 96-well plates were coated with N, E and S SARS-CoV-2 antigens for the evaluation of ADCC functionality. Single donation plasma samples (SARS-CoV-2 negative plasmas) and prepandemic immunoglobulins showed no ADCC activity; on the contrary, IgG hyperimmune samples showed marked ADCC activity for the antigen.

**Figure 2B.**
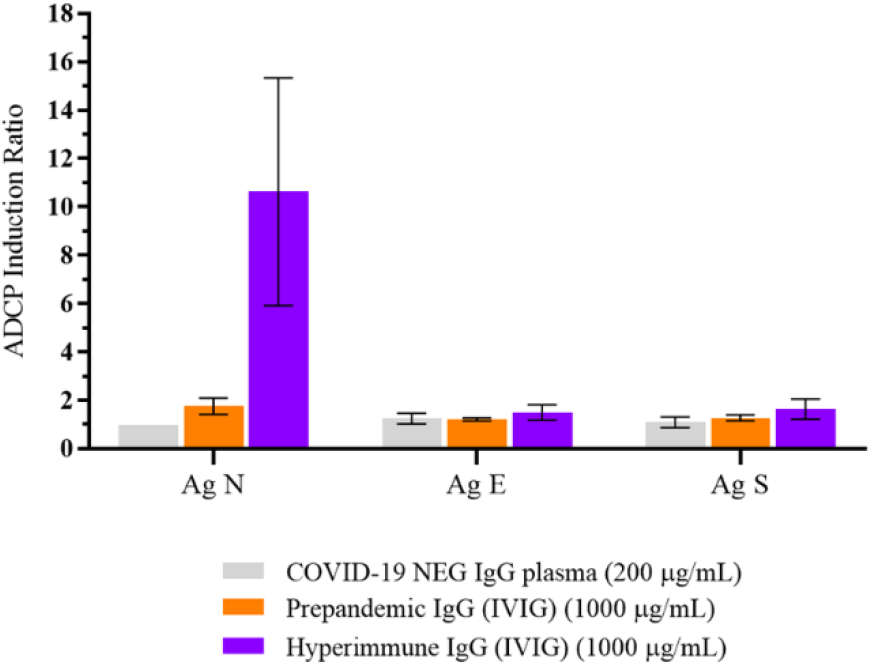
ADCP induction ratio with coated SARS-CoV-2 antigens. 96-well plates were coated with N, E and S SARS-CoV-2 antigens for the evaluation of ADCP functionality. Single donation plasma samples (SARS-CoV-2 negative plasmas) and prepandemic immunoglobulins showed no ADCP activity; on the contrary, IgG hyperimmune samples showed marked ADCP activity for the N antigen.

When several batches of hIG were tested (n= 9), ADCC IR by HEK293T cells expressing SARS-CoV-2 S glycoprotein was above negative control sample IR value for all batches (data analyzed at 100 µg/mL) (Figure 3a). High and medium IgG ELISA SARS-CoV-2 antibody-positive single-donation plasmas were also above the negative control sample IR value, but the low SARS-CoV-2 positive and negative SARS-CoV-2 plasmas were not observed to be above the negative control sample IR. Regarding ADCP IR, the seven hIG batches and the high SARS-CoV-2 positive plasma were above the IR of the negative samples, whereas medium SARS-CoV-2 positive and negative SARS-CoV-2 plasma were not (Figure 3b).

**Figure 3A.**
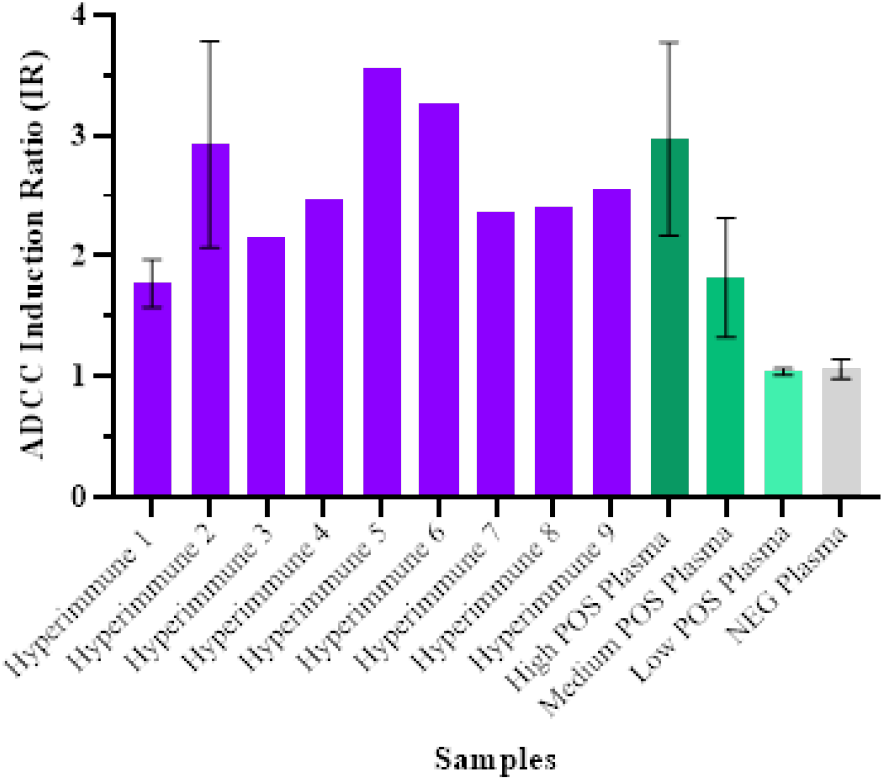
ADCC induction ratio with SARS-CoV-2 Spike Glycoprotein expressing HEK293T cells (Innoprot). Nine hyperimmune samples were assayed and showed high ADCC activity (100 µg/mL) when using S expressing HEK293 cells. SARS-CoV-2 positive plasma samples (100 µg/mL) showed also this functionality.

**Figure 3B.**
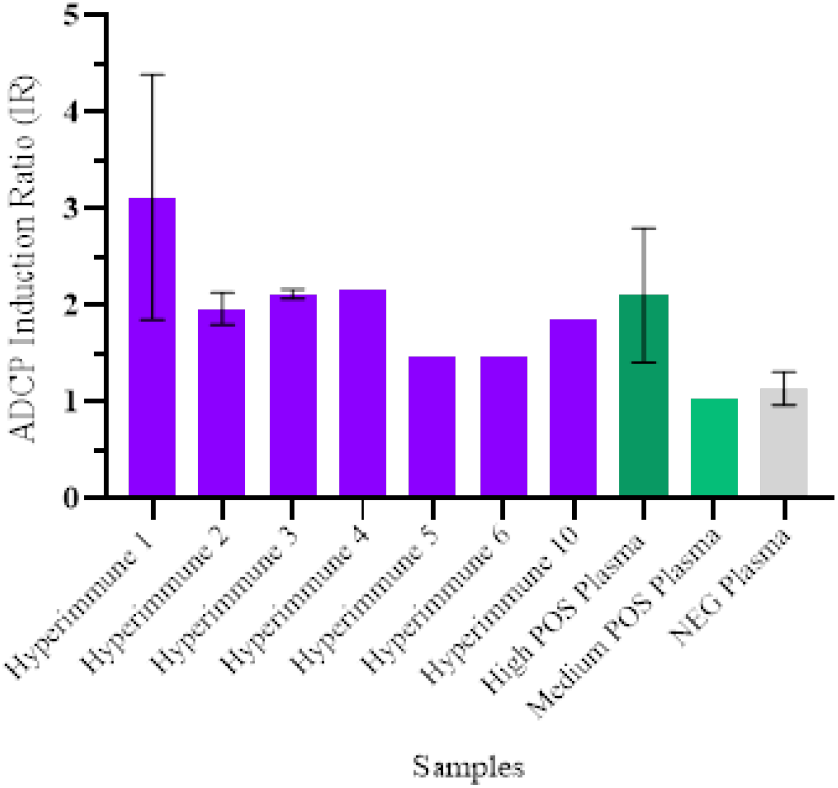
ADCP induction ratio with SARS-CoV-2 Spike Glycoprotein expressing HEK293T cells (Innoprot). Six hyperimmune samples were assayed and showed high ADCC activity (100 µg/mL) when using S expressing HEK293 cells. SARS-CoV-2 positive plasma samples (100 µg/mL) showed also this functionality. Non-statistical differences in ADCP induction ratios among hyperimmune batches are attributable to inter-assay variability.

Both pooled hIG and high SARS-CoV-2 positive plasma showed ADCC induction by HEK293T cells expressing SARS-CoV-2 S glycoprotein. Activity correlated with increasing concentrations. Maximal IR in kinetics studies was observed at 150 µg/mL. Higher concentrations of hIG interfered with the read-out systems (data not shown). Pre-pandemic IVIG samples and other plasmas did not show relevant activity. These data are summarized in Figure 4.

**Figure 4.**
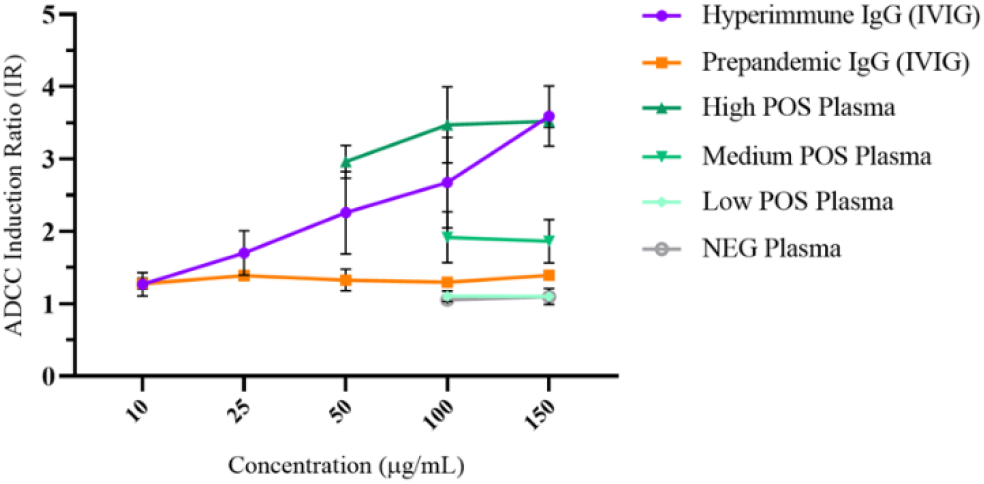
ADCC induction ratio kinetic curve in hyperimmune samples, prepandemic immunoglobulins, SARS-CoV-2 positive (high, medium and low SARS-CoV-2 IgG titers) and SARS-CoV-2 IgG negative single-donation plasmas, in SARS-CoV-2 Spike Glycoprotein expressing HEK293T cells (Innoprot). Hyperimmune batches demonstrate high ADCC functionality at incremental concentrations, with a peak at 150 µg/mL. Non-statistical differences in ADCP induction ratios among hyperimmune batches are attributable to inter-assay variability.

## 4. Discussion

COVID-19 continues to wreak havoc among SARS-CoV-2 infected patients. Despite therapeutic advances aimed at alleviating the symptoms of the disease, no current therapy has been demonstrated to improve outcomes in hospitalized patients in randomized clinical trials [27]. Effective treatments are needed which may more broadly neutralize viral variants through multiple mechanisms.

Here we report an extensive study of the SARS-CoV-2 neutralization capacity of hIG products, not only for different virus isolates from several regions of the world but also for the most relevant SARS-CoV-2 variants studied using a pseudovirus expressing SARS-CoV-2 S glycoprotein. In addition, to our knowledge, this is the first time that several hIG product batches are assayed to evaluate these functionalities showing consistent results. Moreover, the robustness of the anti-SARS-CoV-2 activity of hIG was demonstrated for the first time in different immune effector mechanisms (ADCC, ADCP), with different methodologies, and identifying the viral proteins involved.

All four viral infectivity neutralization methods (CBIFA, CCLA, PFU, TCID50) showed strong neutralization of SARS-CoV-2. ID50 values varied since they are method dependent. Our results using CBIFA were consistent with those reported in the hIG manufacturing characterization (ID50 325±76) [7]. The use of the other neutralization methodologies allowed the detection and reporting of more potent neutralization activity, i.e, higher ID50.

Since high neutralization capacity has been shown employing multiple virus isolates, in several virus-cell systems and with different methodologies, we can describe the neutralization capacity of hIG as robust and likely to be reproducible under normal physiological conditions after administration to patients. The study of the neutralization capacity with a pseudovirus expressing S glycoproteins from the most relevant SARS-CoV-2 variants (second wave variants D614G, B.1.1.7, P.1, and B1.351) is especially important given the current situation in UK, Brazilian and South Africa. In these regions, the predominant variants have been recently classified by the CDC/WHO as variants of great concern [28]. In this study, even if some reduction of pseudovirus neutralization for P.1 and B1.351 has been shown (as preliminary results suggest for some CCP and derived products [29]) effective neutralization of emergent variants is relevant because the plasma used to produce the hIG was collected prior to the detection of these variants. However, these hyperimmune products were demonstrated to have the capability to neutralize these new variants consistently, although this capability has not been observed elsewhere [30].

Beyond neutralization there are other IgG Fc-dependent functionalities that may play a role in the protection from and/or resolution of SARS-CoV-2 infection by hIG products, especially when differences in neutralization activity have been detected for some variants [30, 31].

In antigen-dependent Fc function, studies performed with plates coated with SARS-CoV-2 antigens, only hIG showed relevant ADCC and ADCP activity for the N protein. This is the most abundant protein in coronaviruses. It is highly conserved and is highly immunogenic [32]. This finding could be particularly relevant for variants capable of escaping anti-S neutralization. In fact, S glycoprotein plays a crucial role in viral entry into the cell, which makes it one of the most important targets for COVID-19 vaccine and therapeutic research [33].

However, no apparent activity against E and S proteins was observed in Fc function experiments with antigen-coated plates. While E protein is the smallest of all the structural proteins of SARS-CoV-2, S protein is structurally complex [34]. In both cases, the possibility that the antigen attached to the plate acquired an inadequate conformation to be detected by the test, must be considered. S-protein associated ADCC in COVID-19 patients has been recently reported [35]. Although E protein has recently being considered as a potential therapeutic target [36], we further explored Fc functionality related to the more relevant S protein, using HEK293T cells expressing SARS-CoV-2 S glycoprotein.

In HEK293T S cells, all hIG batches induced considerable ADCC and ADCP activity comparable to that of high titer CCP. This result confirmed that hIG possesses activity against the S protein that was not detected using S antigen-coated plates. Moreover, the ADCC induction ratio was concentration-dependent, with noticeable activity at concentrations as low as 25 µg/mL. At this point, it is important to remember that the IR value of the positive plasma corresponds to a single SARS-CoV-2 positive donor with a high antibody titer, while the hyperimmune IR corresponds to multiple donors with variable positive titers of anti-SARS-CoV-2 immunoglobulins. ADCC and ADCP are mechanisms of action for antigen-dependent antibodies through which virus-infected or otherwise diseased cells are targeted for destruction or elimination. This occurs through multiple components of the cell-mediated immune system, primarily through FcγRIIIa expressed on natural killer cells (for ADCC), and by monocytes-macrophages, neutrophils and dendritic cells via FcγRIIa (CD32a), FcγRI (CD64) and FcγRIIIa (CD16a) for ADCP. The role of antibodies against SARS-CoV-2 N protein in these mechanisms could be a determining factor in resolving SARS-CoV-2 infections through an S-protein independent mechanism. This should be further investigated since, at least theoretically, protein N is not accessible to antibodies in an intact virus or infected cell.

For other viruses such as influenza A, the role of non-neutralizing antibodies against internal and more conserved virus proteins suggest that these antibodies also play an important role in anti-influenza virus immunity [37]. They work by reducing virus titers and ameliorating disease via ADCC [38, 39] and anti-NP antibodies can facilitate particle and antigen uptake and presentation, leading to reduced viral titers and morbidity [39-41].

Monoclonal antibody and vaccine efficacy against SARS-CoV-2 are based on anti-S neutralizing activity. Here we demonstrated the activity of hIG COVID19 through other proteins (N) and through other mechanisms involving host immune system cells (ADCC and ADCP). This opens the door to combine therapeutic and prophylaxis strategies by using these products in combination in order to increase effectiveness.

Knowing the mechanism of action of hIG will help to predict effectiveness and will save time and effort in the selection of the target patient population. In addition, hyperimmune IgG product has anti-viral activities beyond neutralization that when combined with neutralization has the potential to provide a more robust treatment in the future against new infectious threats.

## 5. Conclusions

hIG solutions have strong neutralization capacity against SARS-CoV-2. This strong neutralizing capacity is not only present against viruses that plasma donors were exposed to, but also against the new SARS-CoV-2 emerging variants. In addition, under our experimental conditions, viral N and S proteins induced antigen-dependent Fc functions, such as ADCC and ADCP even at low concentrations. Altogether, this work opens new frontiers in COVID-19 treatment mediated by hyperimmune antibodies. The fact that similar results were obtained with multiple experimental approaches suggest that hIG treatment is a promising therapeutic option for SARS-CoV-2 therapy.

## Acknowledgements

The authors acknowledge the expert technical assistance of the Immunotherapies Unit personnel (D. Casals, J. Luque, E. Sala and M. Blanquer), Bioscience R&D, Scientific Innovation Office (Barcelona, Spain). As well the personnel of the other laboratories that performed the neutralization experiments. Assistance of the Grifols Scientific Publications Department is also acknowledged. E. Pradenas was supported by a doctoral grant from the National Agency for Research and Development of Chile (ANID: 72180406).

## Supplementary Figures

**Supplementary Figure S1.**
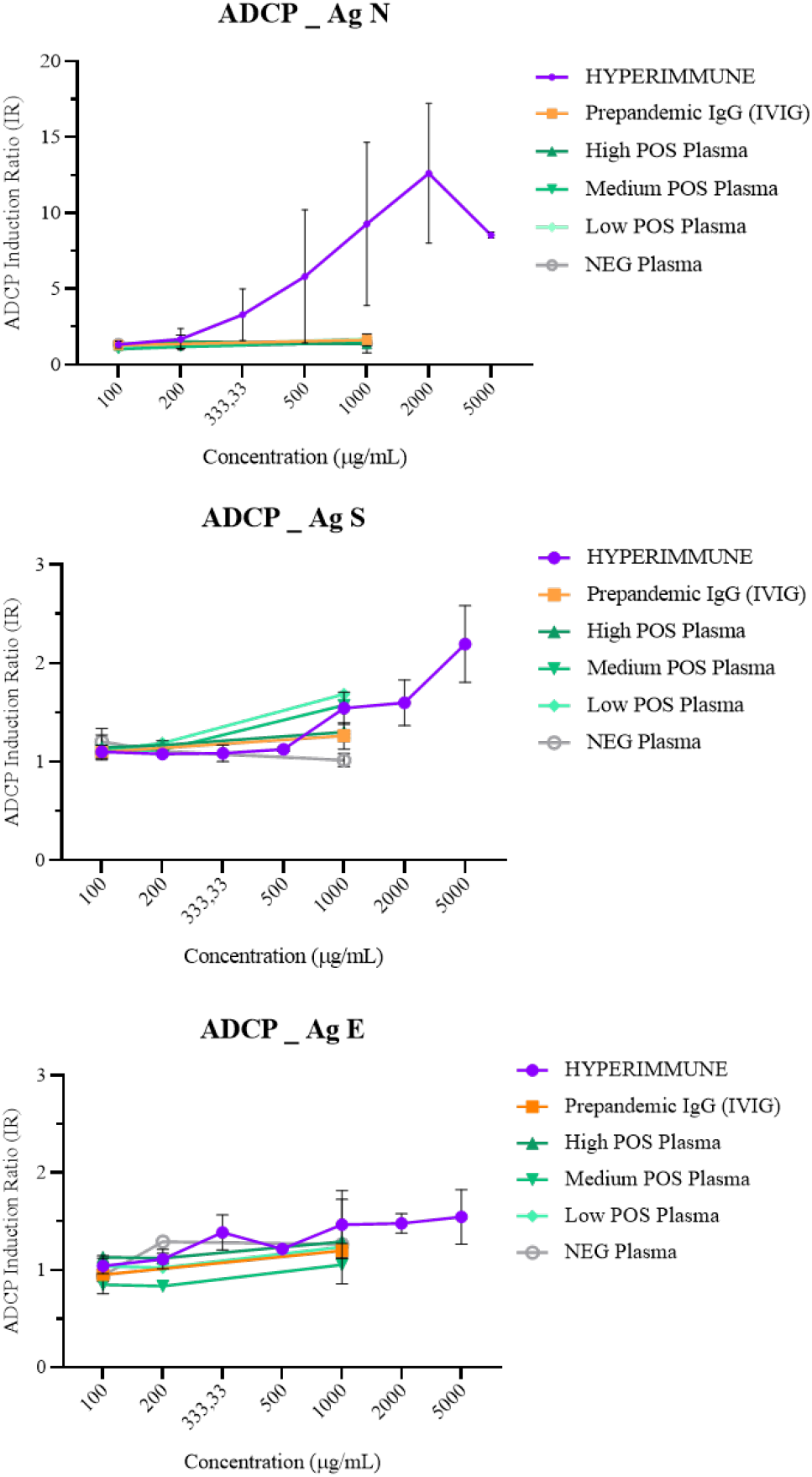
ADCP kinetic curves (µg/mL) in plasma samples (SARS-CoV-2 high, medium and low positive; and SARS-CoV-2 negative single donation plasma samples), prepandemic immunoglobulins and hyperimmune samples. Results show the ADCP induction ratios (IR) obtained at different sample dilutions. Once again, hyperimmune samples showed high ADCP activities at concentrations of 333.33 - 2000 µg/mL with the SARS-CoV-2 N antigen. No positive ADCP functionality is evidenced for any other sample type or SARS-CoV-2 antigen.

**Supplementary Figure S2.**
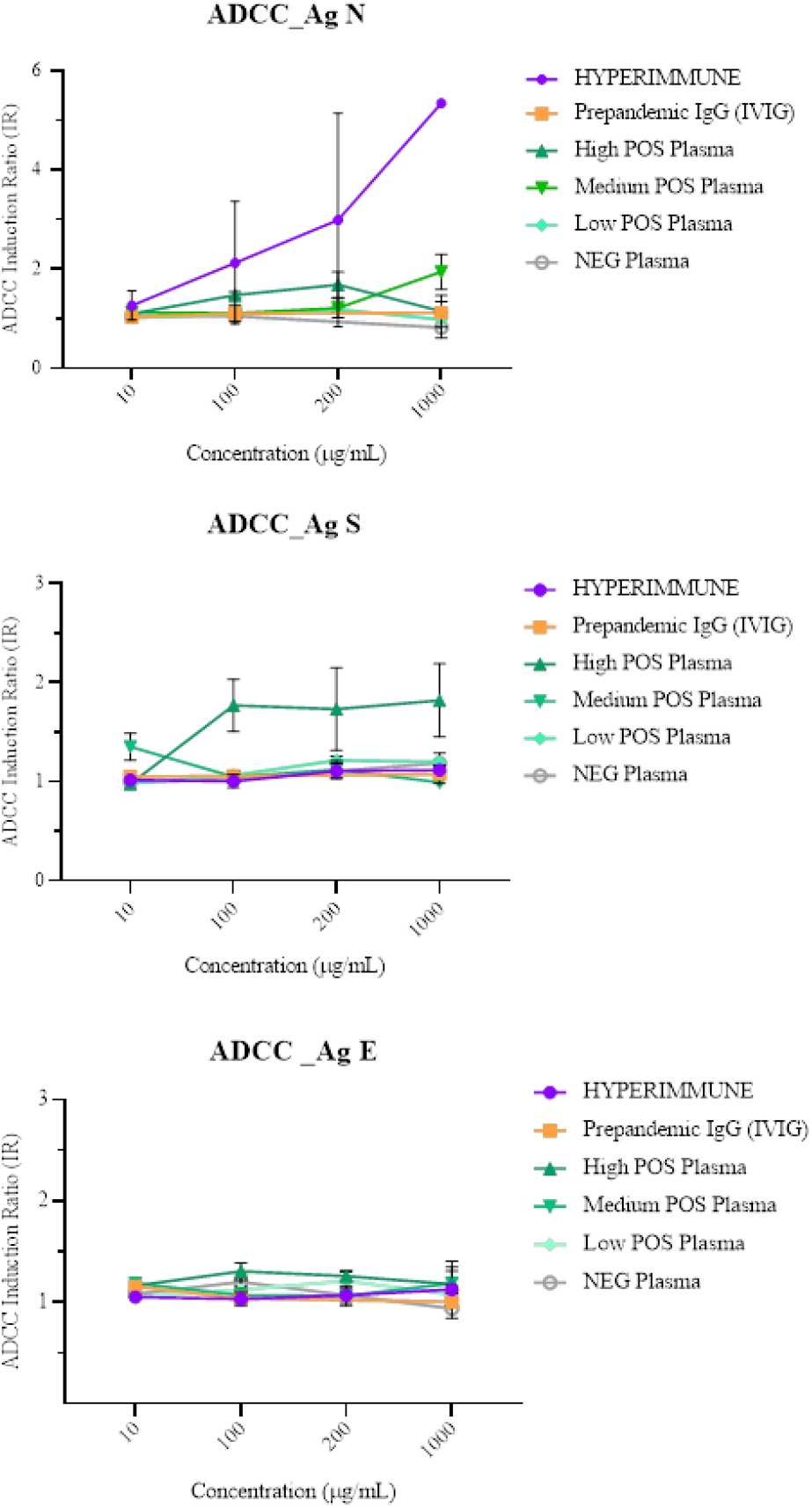
ADCC kinetic curves (µg/mL) in plasma samples (SARS-CoV-2 high, medium and low positive; and SARS-CoV-2 negative single donation plasma samples), prepandemic immunoglobulins and hyperimmune samples. Results show the ADCC induction ratios (IR) obtained at different sample dilutions. Once again, hyperimmune samples showed high ADCC activities at concentrations of 100 - 1000 µg/mL with the SARS-CoV-2 N antigen. No positive ADCC functionality is evidenced for any other sample type or SARS-CoV-2 antigen.

